# Clusters of neuronal Neurofascin prefigure node of Ranvier position along single axons

**DOI:** 10.1101/2021.06.25.449890

**Authors:** Stavros Vagionitis, Franziska Auer, Yan Xiao, Rafael G Almeida, David A Lyons, Tim Czopka

## Abstract

The spacing of nodes of Ranvier crucially affects conduction properties along myelinated axons. It has been assumed that node position is primarily driven by the growth of myelin sheaths. Here, we reveal an additional mechanism of node positioning that is driven by the axon. We show through longitudinal live imaging of node formation dynamics that stable clusters of the cell adhesion molecule Neurofascin A accumulate at specific sites along axons prior to myelination. While some of these clusters change position upon encounter with growing myelin sheaths, others restrict sheath extension and are therefore predictive of future node position. Animals that lack full-length Neurofascin A showed increased internodal distances and less regular spacing of nodes along single axons. Together, our data reveal the existence of an axonal mechanism to position its nodes of Ranvier that does not depend on regulation of myelin sheath length.

## Introduction

The myelination of axons is crucial for fast propagation of action potentials. One of the principal ways in which myelin regulates conduction is by restricting the localization of voltage gated ion channels to discrete axonal subdomains around the nodes of Ranvier – short unmyelinated gaps between each myelin sheath (Rasband and Peles, 2020; Sherman and Brophy, 2005). Not all axons are myelinated equally along their length but can instead show variable and highly organised internodal distances, with implications for conduction speed and circuit function (Brill et al., 1977; Ford et al., 2015; Waxman, 1997; Xin and Chan, 2020). Although we have a detailed understanding of the molecules that organise the architecture of nodes of Ranvier in general (Rasband and Peles, 2020), the mechanisms that determine where nodes are positioned along the length of individual myelinated axons, which ultimately determines their conduction properties, are not understood.

It is generally assumed that the growth of myelin sheaths along axons determines the position of nodes of Ranvier and the distance between consecutive nodes (the internodal spacing). This is due to the fact that a node is characterised as being localised to the gap between two adjacent myelin sheaths, and because the paranodal junctions act as molecular sieves that restrict the localisation of ion channels and adhesion molecules to discrete microdomains around the node (Pedraza et al., 2001). Furthermore, it has been shown that developing myelin sheaths grow to different length in response to neural activity (Baraban et al., 2017; Krasnow et al., 2017). Therefore, the prevailing view is that sheaths push nodes in position and that nodal spacing is thus determined by regulation of myelin growth. However, during the myelination of a single axon along its length, individual sheaths are added at different points in time and by different oligodendrocytes. It remains unclear how specific internodal distancing, such as even spacing along the axon or their progressive shortening towards the synapse, can be achieved by only controlling myelin growth.

One simple alternative mechanism besides oligodendrocyte control of node position would be that it is not only dependent on myelination, but that node position can prefigure myelination and may even restrict myelin sheath growth. One feature of an axon that can inherently limit sheath growth and serve to determine nodal position is their branching pattern. However, we have previously shown that lateral extension of developing myelin sheaths often occurs asymmetrically, even in the absence of obvious physical barriers such as axon collaterals or adjacent myelin sheaths that could restrict lateral myelin growth (Auer et al., 2018). These observations suggest that other axonal cues might restrict sheath growth beyond certain points and therefore determine the position of where a node will form. Interestingly, neurons can cluster sodium channels along unmyelinated axons *in vitro* and *in vivo*, which can even influence conduction speed along such axons (Freeman et al., 2015; Kaplan et al., 1997). However, if the localisation of such pre-myelinating nodal clusters is a determinant of node of Ranvier position along axons *in vivo* is not known.

Here, we investigated the mechanisms of node of Ranvier positioning along single axons using live cell imaging of nodal markers during developmental myelination. We show that the neuronal cell adhesion molecule Neurofascin A (Nfasca) accumulates along single axons prior to myelination. These clusters can either be pushed in place by developing myelin sheaths, or in fact restrict sheath growth and be therefore predictive for future node position. Animals that lack neuronal Nfasca showed increased numbers of very long internodal distances, resulting in less regular nodal spacing along single axons. Therefore, our data reveal the co-existence of two regulatory mechanisms to control node position; one that is intrinsic to the axon and independent of myelin, and another that involves regulation of myelin sheath length.

## Results

### Localisation dynamics of Node of Ranvier reporter constructs during myelination of single axons

In order to investigate mechanisms of node of Ranvier positioning, we studied Commissural Primary Ascending (CoPA) and Circumferential Descending (CiD) neurons of larval zebrafish, which are both glutamatergic spinal interneurons with frequently myelinated axons (Higashijima et al., 2004; Koudelka et al., 2016). Nodes of Ranvier were labelled using three transgenic live cell reporters that localise to nodes by different mechanisms: the GPI-anchored adhesion molecule contactin1a with EYFP fused to its N-terminus (EYFP-cntn1a); the intracellular Ankyrin G targeting motif of zebrafish NaV1.2 (EYFP-NaV-II-III); and neuronal Neurofascin (Nfasca) as a transmembrane protein with a C-terminal fusion of EYFP (Nfasca-EYFP) (Fig. 1A) (Auer et al., 2018; Garrido et al., 2003; Koudelka et al., 2016). Co-expression of these constructs in single neurons alongside a cytoplasmic fluorescent protein to outline axon morphology (cntn1b:XFP) in zebrafish that have all myelin labelled (Tg(mbp:memXFP)) showed that all three nodal reporter constructs specifically accumulate in the gap between two adjacent myelin sheaths (Fig. 1B). EYFP-cntn1a and Nfasca-EYFP fluorescence was completely excluded from myelinated parts of the axon (in agreement with previous observations) (Auer et al., 2018; Koudelka et al., 2016), whereas EYFP-NaV-II-III retained some fluorescence within axon stretches that are ensheathed with myelin (Fig. 1B, B’, D). In contrast to the highly organised localization of fluorescence signal along myelinated and nodal regions, the distribution of all three nodal live cell reporters appeared rather diffuse along unmyelinated stretches of partially myelinated axons (Fig. 1C, S1A-C). However, we did regularly observe accumulations of Nfasca-EYFP fluorescence along unmyelinated stretches that resembled accumulations at nodes and heminodes (Fig. 1C’, S1A). No signs of oligodendrocyte ensheathment could be detected in the vicinity of these Nfasca-EYFP accumulations, nor was Nfasca-EYFP fluorescence excluded along the respective axon stretches as it routinely occurred upon ensheathment, even when myelin sheaths were very short (Fig 1D, S1D). These Nfasca-EYFP clusters were about five times brighter than other unmyelinated regions and of comparable, yet slightly lower, fluorescence intensity than at nodes (rel. Nfasca fluorescence: 0.1±0.03 unmyelinated *vs*. 0.5±0.2 at cluster *vs*. 0.8±0.2 at node, n=7/7 axons/animals, p=0.003 unmyelinated *vs*. cluster, p=0.11 cluster *vs*. node (Kruskal-Wallis test with multiple comparisons); Fig. 1D). We did not detect signs of cluster accumulations using our cntn1a and NaV-II-III reporter constructs (Fig. S1 B, C). The existence of pre-nodal clusters has previously been reported in different systems, including cell cultures and in rodents *in vivo* (Freeman et al., 2015; Kaplan et al., 1997). To confirm that also zebrafish axons can endogenously form such pre-nodal clusters, we carried out whole mount immunohistochemistry for Neurofascin and large calibre axons in a transgenic myelin reporter line. Using this assay, we were able to identify nodes, heminodes, as well Nfasca clusters along unmyelinated axon stretches just as we did using our Nfasca-EYFP reporter (Fig. S1E). Thus, we conclude that Nfasca-EYFP accumulations represent pre-nodal clusters as previously reported in other models.

**Figure 1:**
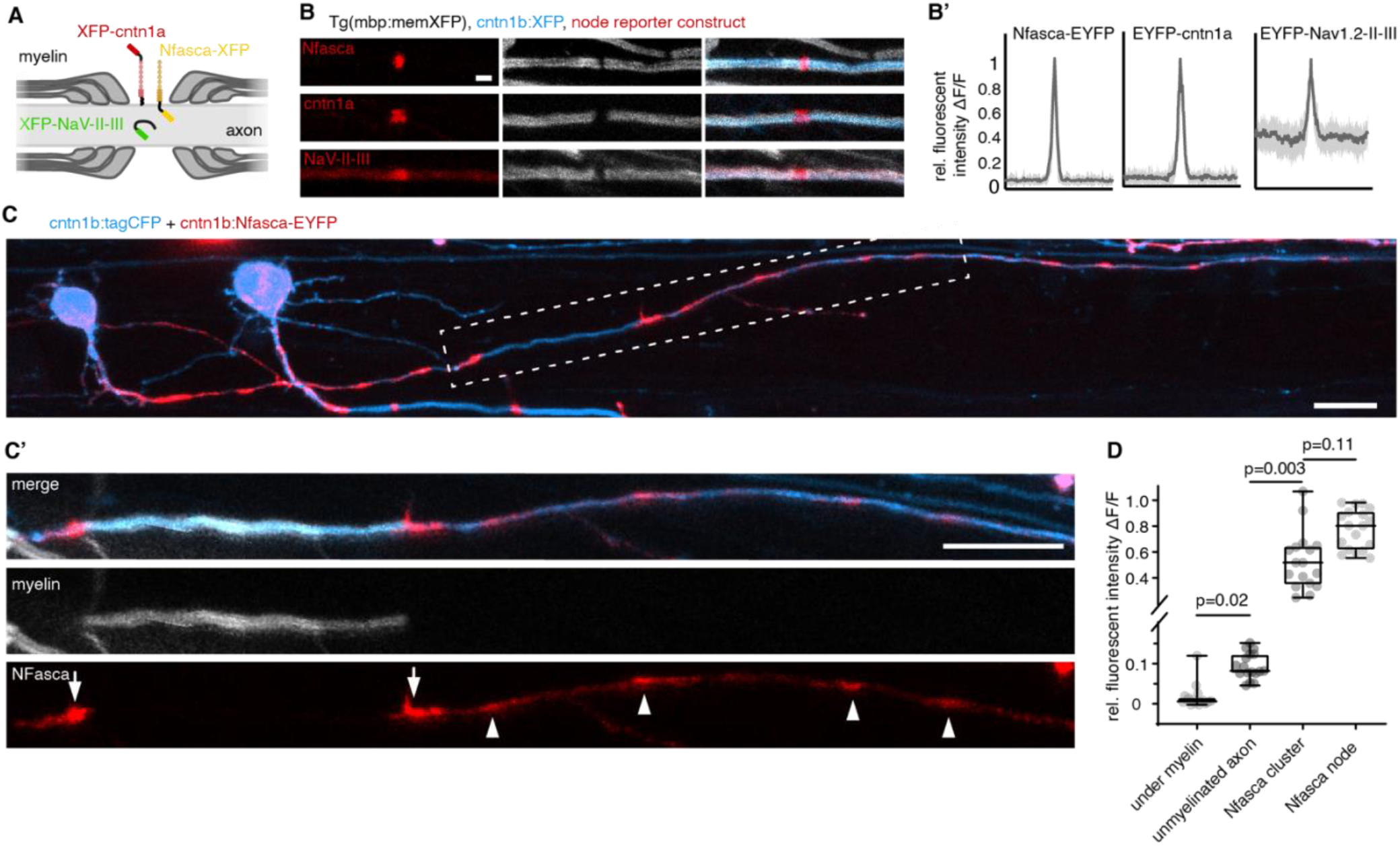
Localisation of transgenic node of Ranvier reporters along unmyelinated and myelinated axon stretches. **A)** Schematic of the node of Ranvier reporter constructs used. **B)** Confocal images of nodal reporter constructs along individually labelled myelinated axons in transgenic animals that have all myelin labelled. Scale bar: 2 μm. **B’**) Fluorescence intensity traces of each reporter construct around the node. Bold lines represent the mean and shaded areas represent SD. **C)** Individually labelled partially myelinated axon at 8dpf co-expressing cntn1b:Nfasca-EYFP in a full transgenic myelin reporter background. **C’**) Magnification of the boxed area in **C**), shown as single channels. Arrowheads indicate accumulations of Nfasca-EYFP along unmyelinated axon parts. Arrows point at Nfasca-EYFP at heminodes. Scale bars: 10μm. **D)** Quantification of fluorescence intensities at regions on analysed axons. Data are expressed as median ± IQR (Kruskal–Wallis test with multiple comparisons). See also Figure S1 for supporting information.

### Neurofascin clusters remain stable along unmyelinated axons and do not associate with processes of oligodendrocyte precursor cells

Having identified pre-nodal clusters in zebrafish, which is an excellent model system for *in vivo* longitudinal imaging over time, we sought to explore the dynamics and fates of these clusters to reveal their role for node of Ranvier positioning during the myelination of individual axons. Consecutive imaging of Nfasca-EYFP labelled single axons revealed that Nfasca clusters along unmyelinated axon stretches were largely stable and comparable to the low motility of bona fide nodes between myelin sheaths (2.5±1.9 μm cluster movement *vs*. 1.8±1.1 μm node movement per day (n=33/5 clusters/axons and 31/9 nodes/axons, p=0.3 (Mann-Whitney test); Fig. 2A, B). Only 6% (4/62) of all clusters analysed disappeared over a two-day period along unmyelinated axon domains (Fig. 2C). It has been reported that processes of oligodendrocyte precursor cells (OPCs) can associate with nodes and pre-nodal clusters to varying extents (Biase et al., 2017; Thetiot et al., 2020). In order to test if OPC processes preferably associate with Nfasca clusters where they could hold them in position, we analysed their association with OPC processes labelled in full transgenic Tg(olig1:mScarlet-CAAX) lines. The vast majority of clusters analysed had no associated OPC processes (60/80), 18/80 clusters had an OPC process crossing in the same imaging plane and only 2/80 clusters were associated with a terminal OPC process (n=4 axons/animals; Fig. 2D-E). Furthermore, these cluster-OPC contacts were only transient and never lasted over the entire period of a 150 minutes timelapse (Fig. 2D’, S2). Therefore, we conclude that Nfasca clusters are stable over time and held in position by a mechanism that is independent of oligodendroglia.

**Figure 2:**
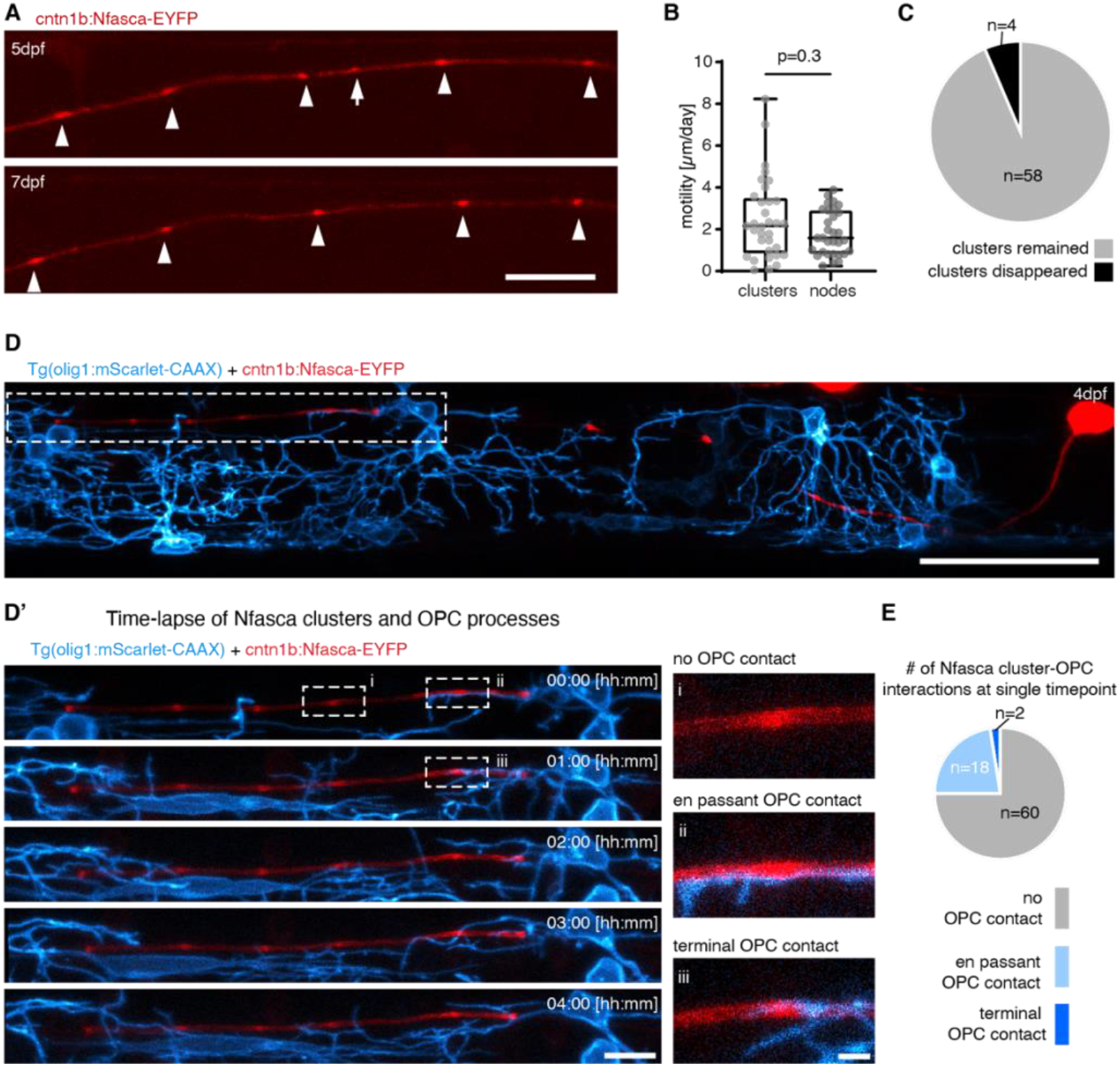
Clusters of neuronal Neurofascin are stable over time and do not co-localise with processes of oligodendrocyte precursor cells. **A)** Confocal image of an individual unmyelinated axon expressing Nfasca-EYFP at two different time points. Arrowheads depict stable clusters and the arrow points to a cluster that disappears between time points. Scale bar: 20 μm. **B)** Quantification of Nfasca cluster and node motility. Data are expressed as median ± IQR (Mann-Whitney test). **C)** Frequency of cluster disappearance. **D)** Confocal image of an axon expressing the Nfasca-EYFP construct in a transgenic animal that has all OPCs labelled. Scale bar: 50 μm. **D’)** (Left) Magnification of the boxed area in panel D showing OPC processes in relation to Nfasca-EYFP accumulations over time. Scale bar: 10 μm. (Right, i-iii) examples of possible cluster-OPC process interactions. Scale bar: 2 μm. **E)** Quantification of the frequency of cluster-OPC process interactions. See also Figure S2 for supporting information.

### Neurofascin clusters can pre-figure node of Ranvier localization

Because clusters of Nfasca fluorescence intersperse axon stretches that are not yet myelinated, we wondered whether these might correlate with future node position. Indeed, Nfasca clusters appeared regularly spaced, but with overall shorter distancing than it was seen for nodes along myelinated axons (18.2±8 μm intercluster distance *vs*. 30.9±13.2μm internodal distance, n=45/6/4 interclusters/axons/animals and 25/4/4 internodes/axons/animals, p<0.001 (unpaired t test); Fig. 3A). These observations are supportive of the idea that at least some Nfasca clusters may form in positions of future nodes of Ranvier. It is known that components of nodes of Ranvier accumulate adjacent to the ends of myelin sheaths to form heminodes (Rasband and Peles, 2020). We found that in our system, 79% of isolated myelin sheath ends that have no neighbouring myelin showed a heminodal Nfasca accumulation (75/95 heminodes, n=22/21 axons/animals) (Fig. 3B). Interestingly, the remaining 21% of myelin sheaths that did not have a Nfasca-labelled heminode were overall shorter than the ones with heminodal Nfasca, both in live cell imaging using our Nfasca reporter as well as in whole mount immunohistochemistry for endogenous Nfasca (Live cell imaging: 35±12 μm *vs*. 23±14 μm, n=58/16 sheaths, p<0.001 (Mann-Whitney test); immunohistochemistry: 25±14 μm *vs*. 11±8 μm, n=72/90 sheaths, p<0.001 (Mann-Whitney test); Fig. 3B, C, S3A).

**Figure 3:**
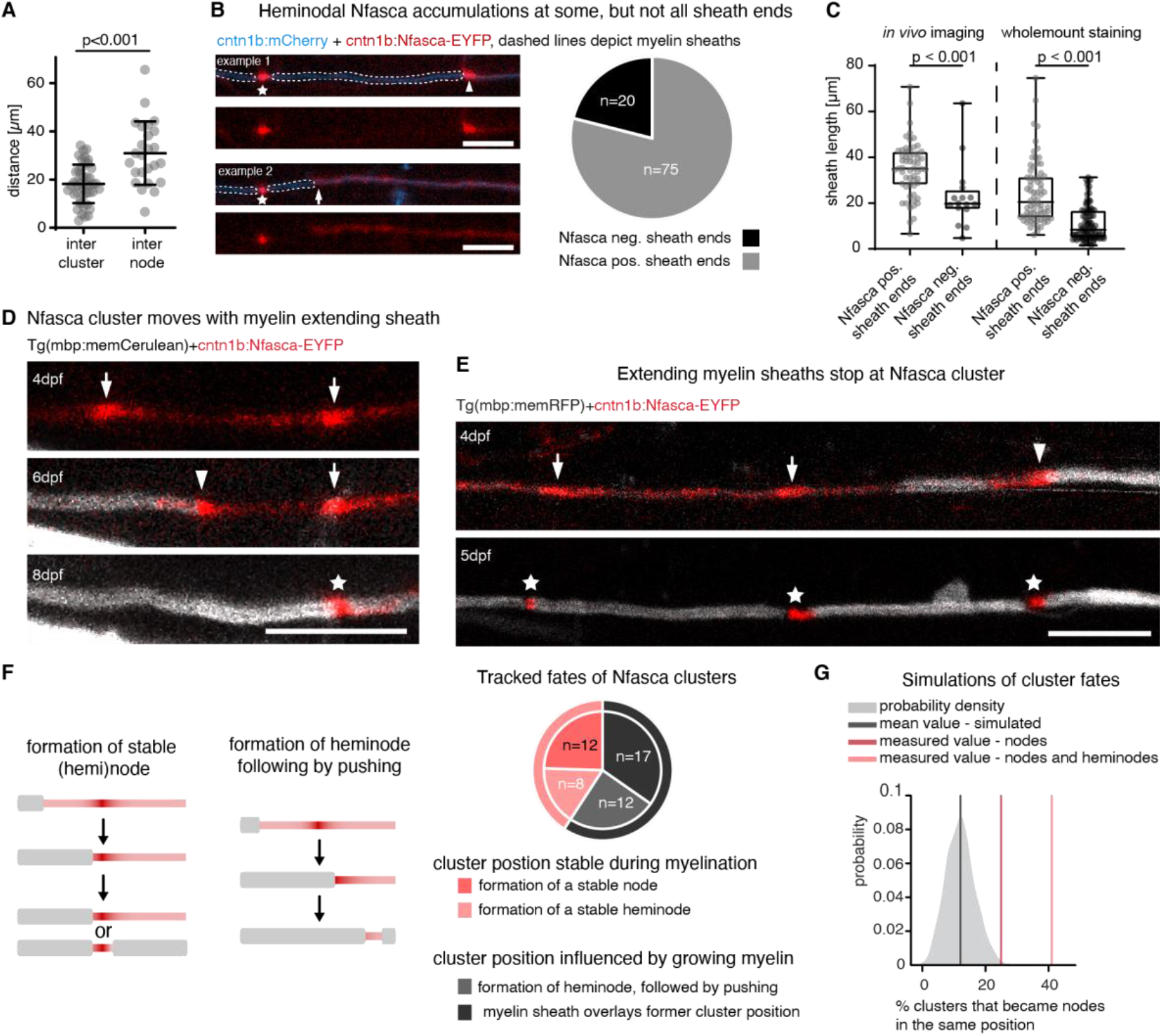
Clusters of neuronal Neurofascin correlate with future node position and restrict myelin sheath extension. **A)** Quantification of intercluster and internodal distances along single axons. Data are expressed as mean ± SD (unpaired t test). **B)** Example images of two individual myelinated areas (dashed lines) forming a node at their left side (asterisk), and the presence (top, arrowhead) and absence (bottom, arrow) of Nfasca-EYFP heminodal accumulation at their right sheath end. Pie chart shows frequency of observations. Scale bar: 10 μm. **C)** Quantification of the length of sheaths with heminodal Nfasca-EYFP compared to sheaths without. Data derived from both *in vivo* imaging (left) and imaging of whole-mount immunohistochemistry (right). Data are expressed as median ± IQR (Mann-Whitney test). **D)** Confocal images of an individual axon expressing Nfasca-EYFP in a full transgenic myelin reporter background at different time points. Arrows depict Nfasca clusters, arrowheads point to heminodes, and asterisks depict nodes of Ranvier. Scale bar: 10 μm. **E)** Confocal images of an individual axon expressing Nfasca-EYFP in a full transgenic myelin reporter background at different time points. Arrows depict Nfasca clusters, arrowheads point to heminodes, and asterisks depict nodes of Ranvier. Scale bar: 10 μm. **F)** Possible cluster fates and quantification of the frequency of events observed by timelapse imaging. **G)** Simulation of the cluster fates and their predicted co-localisation with nodes when random positioning is assumed. The shaded curve shows the probability distribution for the simulated values (grey line indicates the mean of simulated data). The light red and dark red lines indicate the quantified cluster fates analysis shown in panel B. See also Figure S3 for supporting information.

Short myelin sheaths are an indication that they may have formed more recently and are therefore still actively extending in length (Auer et al., 2018). It is conceivable that nascent sheaths could initially form in between clusters, form a heminode when it meets a cluster and continue to grow whilst pushing the cluster until apposing another sheath. In this scenario, sheath growth would determine cluster fate and future node position. Alternatively, developing myelin sheaths stop growing when they meet a cluster. In this scenario, cluster position would pre-determine node position. To test the fates of clusters upon their interaction with developing oligodendrocyte processes, we followed single axons expressing Nfasca-EYFP from unmyelinated through myelinated stages (Fig. S3B). We saw that nascent sheaths could indeed encounter a Nfasca cluster and then continue to grow whilst pushing the cluster by forming a heminode, which ultimately fused with an adjacent myelin sheath to form a node (Fig. 3D). In other cases, however, sheaths did not push the cluster further but instead stopped growing, leading to node formation in this position (Fig. 3E). In order to determine how frequently Nfasca-clusters function as a growth barrier for extending myelin sheaths, which would therefore serve as an indicator of where nodes will form along the length of the axon, we analysed timelines of single axons expressing Nfasca during their myelination between 3 and 8 dpf. Of all 62 clusters analysed (5/5 axons/animals), 9/62 (14.5%) remained clusters because the axon did not get myelinated and 4/62 (6.5%) clusters disappeared or merged with other clusters in the absence of any myelination. Amongst the remaining 49 Nfasca clusters, the position of 40.8% (20/49) was predictive of the location of a node (12/49) or a static heminode (8/49) by the end of analysis, while 59.2% (29/49) of clusters changed in position together with a growing myelin sheath (either by direct observation of heminode formation followed by pushing with growing myelin (12/49) or by a sheath ending up over former cluster position (17/49)) (Fig. 3F). Likewise, a retrospective inspection on nodal origin using all three nodal live cell reporters showed that 55% of all nodes analysed (18/33 nodes in 7 animals) had a Nfasca-EYFP labelled cluster at the same position prior to any sign of myelination (Fig. S3C). No indications for the site of future node positioning prior to myelination were detectable using the nodal reporters EYFP-cntn1a and EYFP-NaV-II-III (Fig. S3C). To rule out that the observed colocalisations between cluster and node position occurred by chance, we simulated how often clusters and nodes would randomly co-localise using the same intercluster and internodal distances observed *in vivo* (Fig. 3A). Our modelling revealed no more than 12±5% probability that a cluster would prefigure node position by chance in a prospective analysis of cluster fates, and no more than 20±7% probability in a retrospective analysis of node origin (Fig. 3G, S3D). In strong contrast, to our measured values showed an at least twofold higher probability that clusters prefigure node position (24.5% (prospective) and 55% (retrospective), and even 40.8% (prospective) and 66.7% (retrospective) when also stable heminodes were included). Together, these data show that Nfasca-EYFP accumulates along axons prior to myelination, and that the positioning of these clusters can correlate with future node of Ranvier position in many cases.

### Increased internodal distances in the absence of full-length Neurofascin A

In order to test if Nfasca itself is functionally important for regulating node positioning, we generated CRISPR mutants of *nfasca*. CRISPR-Nfasca^Δ28^ mutants have a 28bp deletion in the extracellular Mucin domain (specific for the neuronal isoform of Nfasc 186 in mammals), leading to a premature stop prior to the transmembrane domain (Fig. 4A, S4A, B). We confirmed absence of full-length Nfasca protein in homozygous mutants by immunostaining using an antibody against the intracellular C-terminus of Nfasca, which prominently stains nodes in wildtype and heterozygous animals, but not in homozygous mutants (Fig. 4B, B’). In contrast to Neurofascin null mutant mice which are lethal at early postnatal stages (Sherman et al., 2005), our zebrafish *nfasca* mutants are viable through adulthood and appear to breed normally (not shown). *Nfasca* mutants showed grossly normal myelination and even had discrete gaps between adjacent myelin sheaths that accumulated EYFP-cntn1a and EYFP-NaV-II-III reporters (Fig 4C-E), consistent with the premise that myelination with an intact paranodal junction is sufficient to cluster nodal components in the absence of axonal Nfasc 186 in rodents (Zonta et al., 2008). However, analysis of node positioning along single axons using EYFP-cntn1a revealed an overall increase of internodal distances by about 18% in Nfasca mutants over wildtypes (47.1±8.4 in wildtypes *vs*. 56.7±7.7 in *nfasca*^Δ28/Δ28^ in n=10/14 axons 9/14 in animals, p=0.02 (Brown-Forsythe ANOVA with multiple comparisons); Fig. 4F, G). Importantly, increased internodal distances observed in Nfasca mutants could be rescued to wildtype levels by re-expressing Nfasca-EYFP in individual axons of Nfasca mutants (56.7±7.7 in *nfasca*^Δ28/Δ28^ *vs*. 48.5±11.5 in *nfasca*^Δ28/Δ28^ inj. Nfasca-EYFP n=14/14 axons in 14/14 animals, p=0.04 (Brown-Forsythe ANOVA with multiple comparisons); Fig. 4G). Expression of the nodal reporter Nfasca-EYFP and EYFP-cntn1a itself had no effect on internodal distances in wildtype animals (47.1±8.4 for EYFP-cntn1a *vs*. 45.6±10.9 for Nfasca-EYFP in n=10/10 axons in 9/10 animals, p = 0.7 (unpaired t test); Fig. S4D). Closer analysis revealed that this increase in internodal distances seen in Nfasca mutants was caused by an increase in the maximum internode length per axon (83.5±22.4μm in wildtypes *vs*. 110.6±20.4μm in *nfasca*^Δ28/Δ28^ *vs*. 77.8±22.4μm in *nfasca*^Δ28/Δ28^ inj. Nfasca-EYFP; p=0.007 wildtype *vs. nfasca*^Δ28/Δ28^ and p<0.001 *nfasca*^Δ28/Δ28^ *vs. nfasca*^Δ28/Δ28^ inj. Nfasca-EYFP (Brown-Forsythe ANOVA with multiple comparisons); Fig. 4G’, S4C). Minimum internodal length did not change in either condition (21±9.4μm in wildtypes *vs*. 20.8±9.6μm in *nfasca*^Δ28/Δ28^ *vs*. 24±10.1μm *nfasca*^Δ28/Δ28^ inj. Nfasca-EYFP, p>0.7 between all conditions (Kruskal-Wallis test with multiple comparisons, Fig. 4G’’, S4C). As a result, nodes in Nfasca mutants were less regularly spaced and showed an average 44% increase in the range of internodal distances along individual axons, which was again rescued upon re-expression of Nfasca-EYFP in *nfasca*^Δ28/Δ28^ (62.6±21.5μm in wildtypes *vs*. 89.8±25.7μm in *nfasca*^Δ28/Δ28^ *vs*. 53.8±25.4μm *nfasca*^Δ28/Δ28^ inj. Nfasca-EYFP, p=0.005 for wildtype *vs. nfasca*^Δ28/Δ28^ and p=0.001 for *nfasca*^Δ28/Δ28^ *vs. nfasca*^Δ28/Δ28^ inj. Nfasca-EYFP (Brown-Forsythe ANOVA with multiple comparisons); Fig. 4G’”). Therefore, we conclude that axonal Nfasca plays a functional role in regulating the pattern of nodal spacing along individual axons, likely by restricting myelin sheath extension.

**Figure 4:**
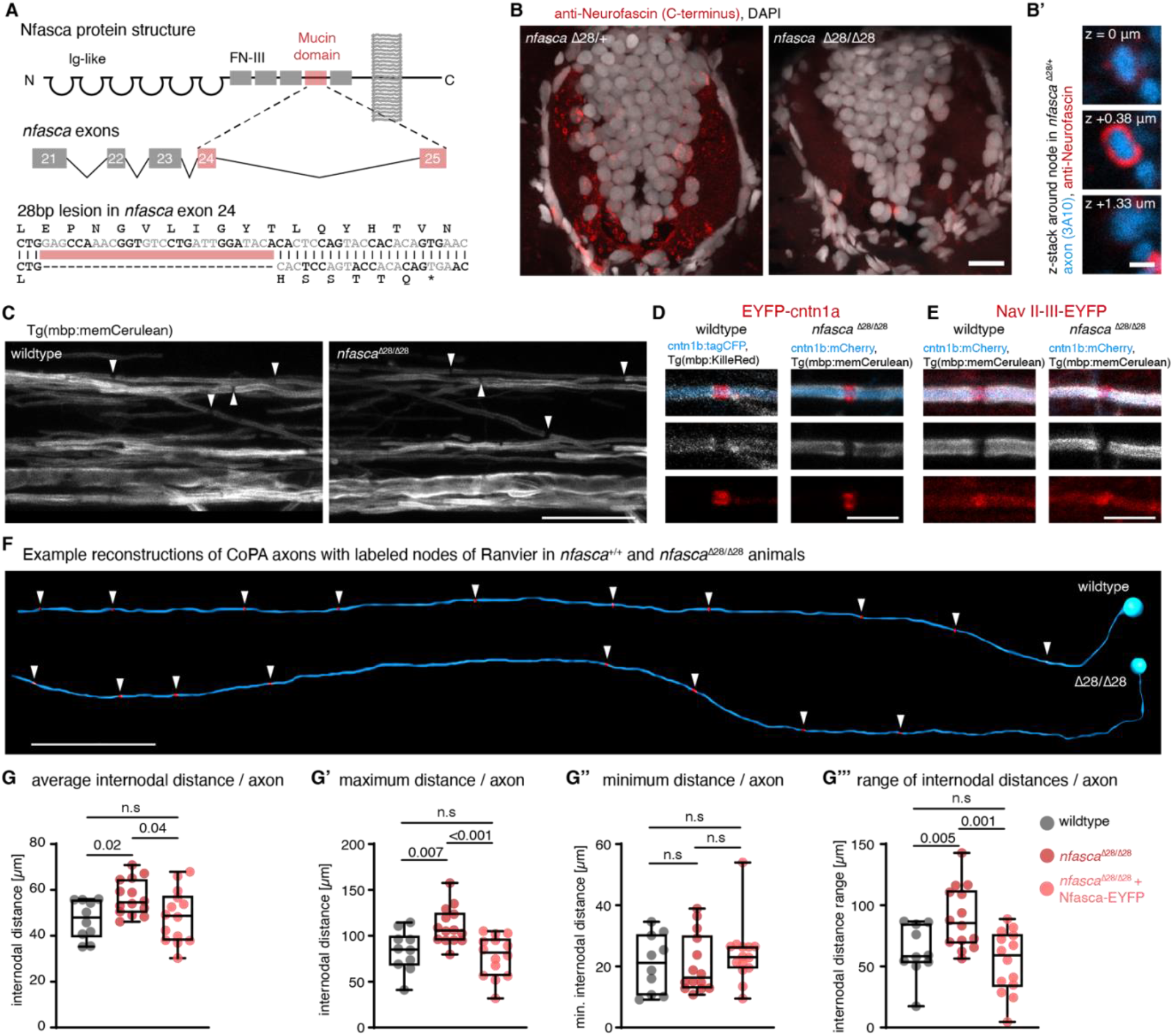
Increased and less regular nodal spacing along individual axons in *nfasca*^Δ28/Δ28^ mutants. **A**) Cartoon illustrating the targeting strategy and genetic lesion in *nfasca*^Δ28/Δ28^ mutants. **B)** Immunohistochemistry against Nfasca on spinal cord cross sections in heterozygous and homozygous *nfasca*^Δ28/Δ28^ animals. Scale bar: 10μm. B’) Cross-sectional views around a node of Ranvier at different z-positions in a heterozygous *nfasca*^Δ28/+^ animal. Scale bar: 1μm. **C)** Confocal images showing overviews of the spinal cord in a full transgenic reporter line for myelin. Arrowheads highlight gaps between adjacent myelin sheaths. Scale bar: 20 μm **D** and **E)** Close-up views showing single nodes of Ranvier marked with EYFP-cntn1a and EYFP-NaV-II-III in a full transgenic reporter line for myelin in WT and *nfasca*^Δ28/Δ28^ mutants. Scale bar: 5 μm. **F)** Reconstructions of two CoPA neurons and their node of Ranvier positions (arrowheads) from wildtype and *nfasca*^Δ28/Δ28^ mutant animals. Scale bar: 50 μm. **G)** Quantification of average internodal distances along single axons per animal in wildtype, *nfasca*^Δ28/Δ28^ and *nfasca*^Δ28/Δ28^ mutants re-expressing Nfasca-EYFP in single axons. Data are expressed as median with IQR (Brown-Forsythe ANOVA with multiple comparisons). **G’-G’’’)**Quantification of maximum (G’), minimum (G’’) and range (G’’’) of measured internodal distances per axon as quantified in G. Data are expressed as median with IQR, (Brown-Forsythe ANOVA with multiple comparisons for G’ and G’’’, Kruskal Wallis test with multiple comparisons for G’’). See also Figure S4 for supporting information.

## Discussion

Our data provide evidence that the axon itself plays an instructive role to determine the position of its nodes of Ranvier. We show that the nodal adhesion molecule Neurofascin A (Nfasca) clusters at specific sites along the axon where they can pre-determine node position by restricting myelin sheath extension. Our work provides novel insights into control of internodal spacing by axons and surrounding oligodendrocytes.

It is well established that formation and maintenance of node of Ranvier architecture requires intrinsic (linkage to axonal cytoskeleton) and extrinsic mechanisms (secreted glial ECM and paranodal adhesion) (Susuki et al., 2013). However, the position where nodes form along an axon had remained essentially uninvestigated. Clustering of sodium channels along unmyelinated axons has previously been observed and it has been speculated that such clusters serve as pre-assembled immature nodes (Freeman et al., 2015; Kaplan et al., 1997). Our data extend current knowledge by providing the first *in vivo* evidence that such pre-myelinating clusters can pre-pattern the axon and therefore determine future node position by restricting sheath growth. Clustering of nodal components requires glial secreted extracellular matrix (Rasband and Peles, 2020). However, axon-intrinsic mechanisms may be sufficient to target components of future nodes at specific sites along the length of axons through their linkage to the axon-cytoskeleton, like Nfasc 186 does by directly binding to Ankyrin G (Davis et al., 1993). Indeed, the evolutionary older axon initial segment (AIS), which shares many features with nodes, does form at a specific site of a proximal axon and does neither require myelin ensheathment nor oligodendrocytes, because invertebrates that have no oligodendrocytes also form an AIS (Hill et al., 2008). Our CRISPR mutants which lack neuronal transmembrane Nfasca show overall increased internodal distances, indicating that Nfasca is either important to form pre-nodal clusters in the first place or to serve as stop signal to prevent myelin sheath extension beyond certain points. We can only speculate why our neuronal Nfasca mutants show no gross neurological defects, which is in contrast to mice that die at early postnatal stages, even upon re-expression of glial Nfasc 155 (Sherman et al., 2005; Zonta et al., 2008). It may be that our mutants still express a functional extracellular portion of Nfasca that is dissociated from the axon membrane, as the stop codon is only just prior to the transmembrane domain. Furthermore, in zebrafish glial and neuronal Neurofascin are encoded by two separate genes (*nfasca* and *nfascb*) (Klingseisen et al., 2019), which may together with the specifics of our CRISPR-Nfasca^Δ28^ gene deletion alleviate the severe phenotypes seen in rodent *nfasc* mutants. Whether CRISPR-Nfasca^Δ28^ affects axons in additional ways than presented here remains to be addressed in future studies.

The presence of axonal landmarks to determine the site of future node position by restricting myelin sheath growth provides a simple mechanism to pattern an axon along its length, a plausible explanation why myelin sheaths along partially myelinated axons are not longer than along fully myelinated axons (Auer et al., 2018; Hill et al., 2018; Hughes et al., 2018), and why developing myelin sheaths mostly grow asymmetrically, despite the absence of obvious physical barriers such as adjacent myelin and axon collateral branches(Auer et al., 2018). It is likely that axonal clustering at future nodal sites is only one of several mechanisms to establish internodal distances along the axon. Observations by us and others argue that either multiple mechanisms are in place at the same time, or that not all axons are pattered by the same mechanisms. For example, it has been reported that only distinct subtypes of axons form clusters (Freeman et al., 2015). Thus, it is possible that pre-nodal clustering to prefigure node position is only employed by subsets of axon types, as it has been shown that neuronal activity promotes myelination of distinct axon types only (Koudelka et al., 2016; Yang et al., 2020). Moreover, our work reveals that not all clusters convert into nodes, but that developing sheaths relocate cluster position and merge several clusters to single nodes, which is in line with another recent study showing that the number of pre-nodal clusters exceeds the number of nodes by about twofold (Thetiot et al., 2020). Our finding that some clusters serve a stop signal for growing sheaths to form a static heminode, while other clusters relocated with the sheath by formation of a migrating heminode argues for the existence of two regulatory mechanisms to control node position; one that is axonal and independent of myelin, and another that involves regulation of myelin sheath length, which can grow to variable lengths depending on the developmental origin of the oligodendrocyte (Bechler et al., 2015), and in response to axonal physiology (Baraban et al., 2017; Krasnow et al., 2017). Future studies will be required to elucidate to which degree these two mechanisms pattern different axon types, and whether one of both of these mechanisms can induce remodelling of internodal distances to adapt axon function.

## Acknowledgements

We are grateful to Wenke Barkey for excellent technical assistance with molecular cloning of the constructs generated for this study and for taking care of the zebrafish colony. We thank Laura Fontenas (Sarah Kucenas group, University of Virginia) for helpful advice with whole mount immunohistochemistry, and all members of the Czopka lab for critical input and discussion of the manuscript. This work was funded by the German Research Foundation DFG (ENP Cz226/1-1 and Cz226/1-2, SFB870 A14 #118803580, and EXC 2145 SyNergy #390857198 to T.C.), a Gertrud Reemtsma PhD student fellowship of the Max-Planck-Society to F.A., support from the TUM PhD program ‘Medical Life Sciences and Technology’ to S.V, and a Medical Research Council (MRC) project grant (MR/P006272/1) to R.A. and D.L.

## Author Contributions

Conceptualisation, F.A., S.V., and T.C.; Methodology, S.V., F.A. and R.A.; Investigation, S.V., F.A., X.Y. and R.A.; Formal analysis, S.V and F.A.; Visualisation, S.V., F.A. and T.C.; Supervision, D.L and T.C; Writing – original draft, T.C.; Funding acquisition, all authors.

## Methods

### CONTACT FOR REAGENT AND RESOURCE SHARING

Further information and requests for resources and reagents should be directed to the lead contact, Tim Czopka and will be fulfilled upon reasonable request. Codes generated during this study are available under https://github.com/Czopka-Lab/Vagionitis-Auer-et-al.

### EXPERIMENTAL MODEL AND SUBJECT DETAILS

#### Zebrafish lines and husbandry

The following existing zebrafish lines and strains were used in this study: Tg(mbp:memCerulean)^tum101Tg^(Auer et al., 2018), Tg(mbp:tagRFPt-CAAX)^tum102Tg^ (Auer et al., 2018), Tg(mbp:KillerRed)^tum103Tg^ (Auer et al., 2018). The following lines have been newly generated for this study: Tg(cntn1b(3kb):KalTA4), Tg(olig1:mScarlet-CAAX), CRISPR-*nfasca*^Δ28^. All animals were kept at 28.5 degrees with a 14/10 hour light/dark cycle according to the local animal welfare regulations. All experiments carried out with zebrafish at protected stages have been approved by the government of Upper Bavaria (animal protocols AZ55.2-1-54-2532-199-2015, ROB-55.2-2532.Vet_02-15-200, and ROB-55.2-2532.Vet_02-18-153 to T.C.).

### METHOD DETAILS

#### Transgenesis constructs

The following existing expression constructs were used in this study: pTol2_cntn1b:Nfasca-EYFP (Auer et al., 2018), pTol2_cntn1b:mCherry (Czopka et al., 2013). The following expression constructs and entry clones were newly generated for this study, with the sequences of all primers used listed in Table S1: p5E_cntn1b(3kb), for which we cloned a 3kb fragment upstream of the start codon and until the second exon of the *cntn1b* gene (ENSDARG00000045685) from genomic DNA of AB zebrafish. The PCR product was recombined with pDONRP4-P1R using BP clonase (Invitrogen). pME_EYFP-NaV-II-III, for which a 1.3 kb fragment corresponding to the II-III region of Nav1.2 (Garrido et al., 2003) of the zebrafish *scn1lab* gene (between exons 16 and 18 of ENSDART00000151247.3) was commercially synthesized (BioCat). We then PCR amplified NaV-II-III from this donor plasmid using XhoI-NaV-II-III_F and XbaI-NaV-II-III_R primers and used these sites to clone it into pCS2+ containing EYFP with the stop codon removed between BamHI and EcoRI sites. The resulting plasmid pCS2+_EYFP-NaV-II-III was then used as template for a PCR reaction using attB1_YFP_F and attB2R-NavNaV-II-III_R primers, and recombination cloned into pDONR221 using BP clonase, creating pME_EYFP-NaV-II-III. To generate pME_EYFP-cntn1a, we first amplified 2.6 kb of the zebrafish *contactin1a* gene (Ensembl: ENSDART00000170348.2) lacking base pairs 1-135 (amino acids 1-45 coding for signal peptide for secretory pathway) from zebrafish cDNA using the primers EcoRI_cntn1a_F and XbaI_cntn1a_R and cloned into pCS2+ containing EYFP fused to the secretory pathway signal peptide of zebrafish *cd59* (ENSDART00000126737.4) between BamHI and EcoRI sites to create pCS2+_sigpepEYFP-cntn1a. This plasmid was used as PCR template using the primers attB1_sigpepEYFP_F and attB2R_cntn1a_R and the PCR product was recombination cloned in pDONR221 using BP clonase to generate pME_EYFP-cntn1a. To generate pME_mScarlet-CAAX we used the commercially synthesized construct pUC57_attB1_mScarlet_attB2R (BioCat). This plasmid was then used as template for a PCR reaction using attB1_F and CAAX-mScarlet_R. The product of this PCR was used as a template for a subsequent PCR reaction with the primers attB1_F and AttB2R-CAAX-R and the product of this reaction was cloned into pDONR221 using BP clonase, creating pME_mScarlet-CAAX. The expression constructs pTol2_cntn1b(3kb):KalTA4, pTol2_cntn1b(5kb):EYFP-cntn1a, pTol2_cntn1b(5kb):EYFP-NaV-II-III, pTol2_UAS(10x):EYFP-cntn1a, pTol2_cntn1b(5kb):tagCFP, and pTol2_olig1:mScarlet-CAAX were created via multisite LR recombination reactions using the newly generated entry clones described above, in addition to p5E_cntn1b(5kb) (Czopka et al., 2013), p5E_olig1 (Marisca et al., 2020), pME:KalTA4 (Almeida and Lyons, 2015), pME:tagCFP (Auer et al., 2018) and p5E_UAS, p3E-pA, and pDestTol2_pA of the Tol2Kit (Kwan et al., 2007).

#### DNA microinjection and generation of transgenic lines

Zebrafish eggs were injected at the one cell stage using a pressure microinjector with 1nl of injection solution (20-30 ng/μl plasmid DNA, 25-60 ng/μl transposase mRNA and 1% phenol-red (Sigma Aldrich)). Injected animals were screened under a stereo-dissecting fluorescence microscope at 3-4 dpf for transgene expression and used for subsequent in vivo microscopy or were raised to adulthood. Injected adults were outcrossed to wild type animals and their progenies were screened for germline transmission to raise stable transgenic lines.

#### Generation of nfasca mutants using CRISPR/Cas9

Zebrafish CRISPR-Nfasca^Δ28^ mutants were generated using CRISPR/Cas9 genome editing by injecting 12.5 ng/μl sgRNA and 300 ng/μl Cas9 encoding mRNA in AB wildtype embryos at the 1-cell stage. The sgRNA against exon 24 of the *nfasca* gene was designed using CHOPCHOP (Labun et al., 2016) and generated as described (Hruscha et al., 2013). F1 animals were sequenced for indel mutations and an individual with a 28 bp deletion 5’ to the target site was selected to raise CRISPR-Nfasca^Δ28^.

#### Mounting of zebrafish for live cell microscopy

For in vivo cell imaging, we anaesthetized zebrafish embryos and larvae using 0.2 mg/ml MS-222 (PHARMAQ, UK). Animals were mounted laterally in 1% ultrapure low melting point agarose (Invitrogen) on a grass coverslip as previously described in detail by us (Vagionitis and Czopka, 2018).

#### Immunohistochemistry

Zebrafish embryos or larvae were euthanised with an overdose of MS-222 (15 mM) and fixed overnight in 4% paraformaldehyde at 4°C. Following washes in phosphate buffer saline (PBS- 3 × 10 min), fish were incubated in incrementally increasing concentration of sucrose (10, 20 and 30%) until full submersion. Animals were mounted in TissueTek (OCT), sectioned with 14-17μm thickness on a cryotome (Leica CM1850 UV) and subsequently stored at −20°C until further use. For immunofluorescence staining, sections were rehydrated using PBS and subsequently boiled in 10mM citrate buffer, pH=6 for 10 min for antigen retrieval. After 30 min of cooling down in RT and subsequent PBS washes, the slides were permeabilised with PBS containing 0.1% Tween-20 (PBST) for 10-15 minutes. Following blocking in PBST containing 10% fetal calf serum (FCS), 3% normal goat serum (NGS) and 1% bovine serum albumin (BSA), slides were incubated with the primary antibodies in blocking solution overnight at 4°C, washed three times in PBST, and incubated with the secondary antibodies for 1h at room temperature. Finally, slides were washed two times with PBST, once with PBS and then mounted under a coverslip using Prolong Gold Antifade containing DAPI (Invitrogen).

#### Whole-mount Immunohistochemistry

Zebrafish embryos or larvae were euthanised with an overdose of MS-222 (15 mM) and fixed in 2% paraformaldehyde and 1% trichloroacetic acid for 10 minutes at room temperature. Samples were washed in phosphate buffer saline supplemented with 0.1% Tween20 (PBST- 3 x 5 min). Subsequently, fish were incubated in distilled water for 5 minutes at room temperature. Animals were then permeabilized by being incubated for 10 minutes in acetone at −20 °C. A second wash in distilled water was followed by a wash in PBST, both for 5 minutes at room temperature. Blocking was the same as described in the previous paragraph. Primary antibody incubation lasted for 68-72 hours at 4 °C. This was followed by 3 washes of 1 hour each with PBST at 4 °C. Then the samples were incubated with the secondary antibodies overnight at 4 °C. Finally, the fish were again washed 3 times for at least 1 hour each at 4 °C, followed by a final wash with PBS for 15 minutes at 4 °C and embedding in 1% LMP agarose for confocal microscopy.

#### Microscopy and image acquisition

For imaging of embedded zebrafish embryos and larvae we used a confocal laser scanning microscope (Leica TCS SP8). Individual axons were identified and analysed along the whole length of the spinal cord (soma location somite level 5 to somite level 27). DAPI was excited with 405 nm wavelength, Cerulean and tagCFP with 448 and 458nm, ALEXA488 with 488nm, EYFP with 514nm, tagRFPt, mCherry, and ALEXA555 with 552 and 561nm, and ALEXA633 with 633nm. During in vivo microscopy, confocal z-stacks were acquired as 12bit images with a pixel size ranging between 76 and 114 nm and a z-spacing of 1μm using a Fluotar VISIR 25x/0.95 WATER immersion objective. Images are shown as lateral views of the zebrafish spinal cord with the anterior to the left and dorsal to the top of the image. Confocal z-stacks of zebrafish spinal cord cross-sections were acquired as 12bit images with a pixel size of 36 nm, z-spacing of 0.19 μm and pinhole set to 0.8AU using an HC PL APO CS2 63x/1.20 WATER immersion objective.

### QUANTIFICATION AND STATISTICAL ANALYSIS

#### Reconstructions of confocal images

We used the FilamentTracer tool of Imaris 8.4.2 (Bitplane) to reconstruct the axon morphologies and node positioning. The cytoplasmic marker expressed by the axon was used for manual axon tracing while the nodal marker was automatically traced. The two traces were superimposed for visualisation of the node spacing along single axons.

#### Fluorescence intensity measurements

Fluorescent intensities were measured using the x line tool in Fiji (distribution of ImageJ) with a line width adjusted to the thickness of each axon. For each axon, fluorescence intensity values were manually allocated to node, cluster and myelinated categories. One value for the background was obtained for each image and was subtracted. For the intensities of markers at the nodes of Ranvier, 1μm-long axon stretches were measured and averaged. Several nodes and myelinated areas were measured. The fluorescence intensity traces were normalised to the maximum value of the node and the peak was aligned. Average and standard deviation for all nodes were calculated and plotted.

For the partially myelinated axons, one node and one myelinated stretch were measured as well as the unmyelinated part, and all values were normalised to the maximum intensity of the brightest node. For the fluorescent intensity traces, the values were again normalised to the average fluorescence of 5μm of myelinated axon. ΔF/F was calculated and plotted.

For the swarmplot, background was subtracted. For comparing fluorescent intensities at different positions along the axon, a Kruskal-Wallis test with Dunn’s correction for multiple testing was used.

#### Node cluster correlation

Clusters were detected by visual impression, as obviously denser accumulations of nodal reporter construct compared to the diffused localisation along unmyelinated parts of the axon devoid of clusters. For the retrospective analysis, images were aligned to a landmark close to the node analysed. If more than one node was analysed on the same axon, the images were realigned for each node. The growth factor of the animal between two imaging sessions was determined as the % change in linear distance between 2 neuronal somata in the spinal cord. This factor was used to correct the aligned images for body growth. A box with width equal to node width (1-2 μm) was drawn and superimposed to the aligned images from earlier timepoints. Only when a cluster or a heminode was found inside this border, it was counted as residing at the same position.

For the prospective analysis, images were again aligned to a landmark close to the analysed cluster. Again, alignment was corrected for body growth. A box of 2-3 μm width was drawn around each cluster and it was used to assess the cluster fate over time. In these analyses, only clusters and nodes devoid of axon collateral branches were analysed, to exclude the effect of structural obstacles in the analysis.

#### Simulation of node cluster correlation

Cluster to node correlation was simulated using a custom written Matlab script available on https://github.com/Czopka-Lab/Vagionitis-Auer-et-al. The intercluster/-node distances were randomly generated from normal distributions with the observed means and SDs. Cluster and node width was set to 2μm The same number of cluster and nodes, respectively, were generated as in the analysed data. From the intercluster/-node distances the position on the simulated axons are calculated by adding them and also adding the cluster/node width after each intercluster/-node distance.

For the prospective analysis for each cluster, it was checked if there was a node at the same position. Therefore, the positions of the node and cluster were subtracted, and if the absolute difference was less than the node width it was counted as colocalisation. For the retrospective analysis, the same analysis was carried out but for each node it was checked if there was a cluster at the same position. The models gives the mean percentage and SD of by-chance colocalisations as well as the maximum observed values for both analyses.

#### Sheath length measurements

Individual isolated sheaths of partially myelinated axons were visually selected for being flanked by one or two heminodes. Sheath length was measured using the x line tool of ImageJ. Sheaths were then categorised based on the presence or absence of Nfasca accumulations at their ends.

#### Internodal and intercluster distance measurements

The distance along the axon between discrete nodal marker accumulations (clusters or nodes) was measured using the x line tool of ImageJ. The distance between the soma and the first node/cluster was not measured.

For the CRISPR-Nfasca^Δ28^ analysis, internodal distance was measured along CoPA axons in the spinal cord of 8 and 9 dpf zebrafish. Average internodal distance was calculated for each animal and was directly compared to WT. Analysis was blind, as microscopy and analysis was completed prior to genotyping. Average, range, minimum and maximum internodal distance per individual animal was then plotted.

#### Node and cluster dynamics

The distance along individual axons between a node and a chosen landmark was measured over time. The measured distance was corrected for body growth by dividing by the respective growth factor, which was obtained by measuring the distance between two landmarks over time. The corrected internodal lengths were subtracted to calculate daily motility. For the node translocation plot, the initial position of the node was set to 0.

The directionality of movement was not analysed and all movements are expressed as absolute values.

#### Nfasca cluster and OPC process interactions

Single images of axons expressing Nfasca-EYFP and showing clusters were analyzed for the presence or not of an OPC process in contact with the clusters on the same 1 μm z-plane. Found contacts were categorized as en passant, when the OPC process seemed to pass by a Nfasca-EYFP cluster or terminal, when the OPC process tip was in contact with the observed cluster.

#### Statistics

Mean and standard deviation were always calculated using GraphPad Prism9 (GraphPad Software LCC) descriptive statistics tool. Normality tests were used to assess the Gaussian distribution of the data. Normally distributed data are expressed as mean with standard deviation (SD). Datasets that contain non-normally distributed data are expressed as median with interquartile range (IQR). Statistical testing was done appropriately according to the type of distribution of data. For simplicity and legibility, the main text always reports mean and standard deviation. Differences in the fluorescence intensity measurements were tested using the Kruskal-Wallis test with multiple comparisons, controlling for False Discovery Rate (FDR) with the two-stage step-up method of Benjamini, Krieger and Yekutieli. Differences in cluster and node motility were tested using the Mann-Whitney test, as was the difference between lengths of sheaths that showed Nfasca-positive or Nfasca-negative heminodes. Differences between intercluster and internode distance were tested using an unpaired t-test. Differences in average internodal distances between WT, Nfasca mutant and rescued Nfasca-EYFP expressing mutant were tested using Brown-Forsythe ANOVA with multiple comparisons. Differences in range of internodes measured per axon between WT, Nfasca mutant and rescued Nfasca-EYFP expressing mutant were tested using Brown-Forsythe ANOVA test with multiple comparisons, as were the differences in maximum measured values per axon. Differences in minimum of internodes measured per axon between WT, Nfasca mutants and rescue Nfasca-EYFP expressing mutants were tested using Kruskal-Wallis test with multiple comparisons. All multiple comparisons tests in the analyses of WT, Nfasca mutant and rescued Nfasca-EYFP expressing mutant were performed controlling for False Discovery Rate (FDR) with the two-stage step-up method of Benjamini, Krieger and Yekutieli.

### KEY RESOURCES TABLE

| REAGENT or RESOURCE | SOURCE | IDENTIFIER |
| --- | --- | --- |
| *Antibodies* |  |  |
| chicken anti-GFP | Abcam plc. | ab13970 |
| mouse IgG1 anti-3A10 | DSHB | AB_531874 |
| rabbit anti-Neurofascin | Abcam plc. | ab183082 |
| goat anti-mouse IgG, Alexa Fluor 663 | Thermo Fisher | A-21052 |
| goat anti-rabbit IgG, Alexa Fluor 555 | Thermo Fisher | A-21428 |
| goat anti-chicken IgY, Alexa Fluor 488 | Thermo Fisher | A-11039 |
| *Experimental Models: Organisms/Strains* |  |  |
| Tg(mbp:memCerulean)<sup>tum101Tg</sup> | Auer, 2018 | ZFIN: ZDB-ALT-180712-3 |
| Tg(mbp:KillerRed)<sup>tum103Tg</sup> | Auer, 2018 | ZFIN: ZDB-ALT-180712-5 |
| Tg(mbp:tagRFPt-CAAX)<sup>tum102Tg</sup> | Auer, 2018 | ZFIN: ZDB-ALT-180712-4 |
| Tg(cntn1b(3kb):KalTA4) | This paper | N/A |
| Tg(olig1:mScarlet-CAAX) | This paper | N/A |
| CRISPR-Nfasca<sup>Δ28</sup> | This paper | N/A |
| *Oligonucleotides* |  |  |
| For all oligonucleotides used, see Table S1 | this paper | N/A |
| *Recombinant DNA* |  |  |
| p5E_cntn1b(3kb) | this paper | N/A |
| pME_EYFPnostop_NaV-II-III | this paper | N/A |
| pME_sigpepEYFP_cntn1a | this paper | N/A |
| p5E_cntn1b(5kb) | Czopka, 2013 | N/A |
| pME_KalTA4GI | Almeida & Lyons, 2015 | N/A |
| p5E_UAS(10x) | Kwan, 2007 | N/A |
| p3E_pA | Kwan, 2007 | N/A |
| pDestTol2_pA | Kwan, 2007 | N/A |
| pME:tagCFP | Auer, 2018 | N/A |
| pTol2_cntn1b(5kb):Nfasca-EYFP | Auer, 2018 | N/A |
| pTol2_cntn1b(5kb):EYFP-cntn1a | this paper | N/A |
| pTol2_cntn1b(5kb):EYFP-NaV-II-III | this paper | N/A |
| pTol2_UAS(10x):EYFP-cntn1a | this paper | N/A |
| pTol2_cntn1b(5kb):tagCFP | this paper | N/A |
| pTol2_cntn1b(5kb):mCherry | Czopka, 2013 | N/A |
| *Software and Algorithms* |  |  |
| Fiji | http://fiji.sc/ | RRID:SCR_002285 |
| Imaris | Bitplane | RRID:SCR_007370 |
| Imaris FilamentTracer | Bitplane | RRID:SCR_007366 |
| Huygens Essential | Scientific Volume Imaging | RRID:SCR_014237 |
| Graphpad Prism 8 | Graphpad Software | RRID:SCR_015807 |

**Figure S1:**
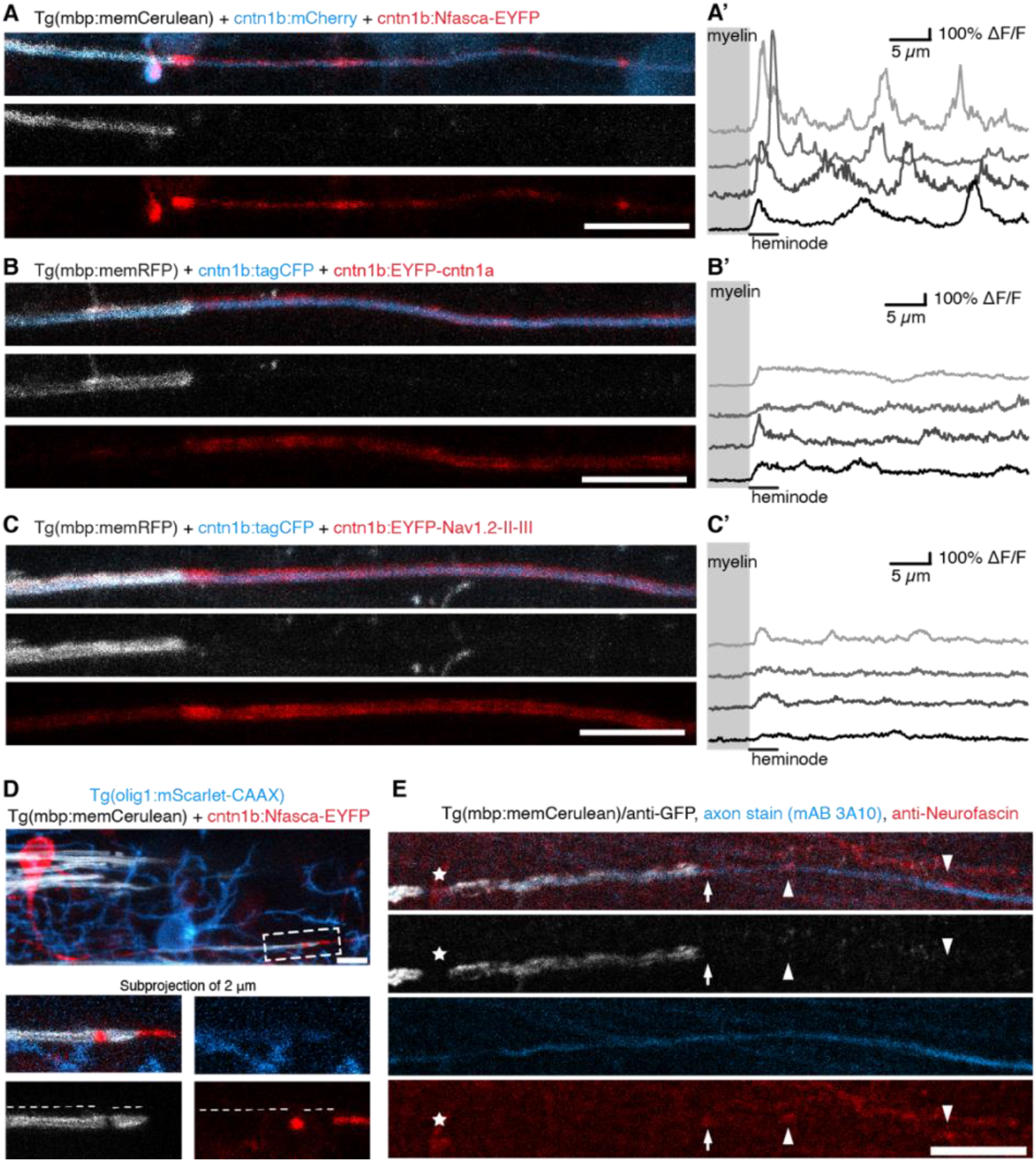
Supplementary to Figure 1. **A-C)** Confocal images of a partially myelinated axon expressing cntn1b:Nfasca-EYFP (A), cntn1b:EYFP-cntn1a (B), and cntn1b:NaV-II-III-EYFP (C). Scale bars: 10 μm. A’-C’ show respective fluorescence intensity profiles along four partially myelinated axon stretches. Grey box indicates myelin sheath position. **D)** Confocal image of a single neuron expressing cntn1b:Nfasca-EYFP in a full-transgenic animal labelling OPCs and myelinating oligodendrocytes. Boxed area in left image shows the area around a node with one established (left) and one very short myelin sheath (right). Single channels reveal complete exclusion of Nfasca fluorescence even under very short myelin sheaths (dashed lines). Scale bar: 10 μm. **E)** Confocal images of a whole-mount immunohistochemistry staining against 3A10 (large calibre axons – cyan) and Nfasca (red) in the spinal cord of a full transgenic line with all myelin labelled (+staining against GFP – grey). Arrowheads indicate a Nfasca cluster along an unmyelinated axon part. Arrows indicates a heminode and the asterisk indicates the presence of a node of Ranvier. Scale bar: 10 μm.

**Figure S2:**
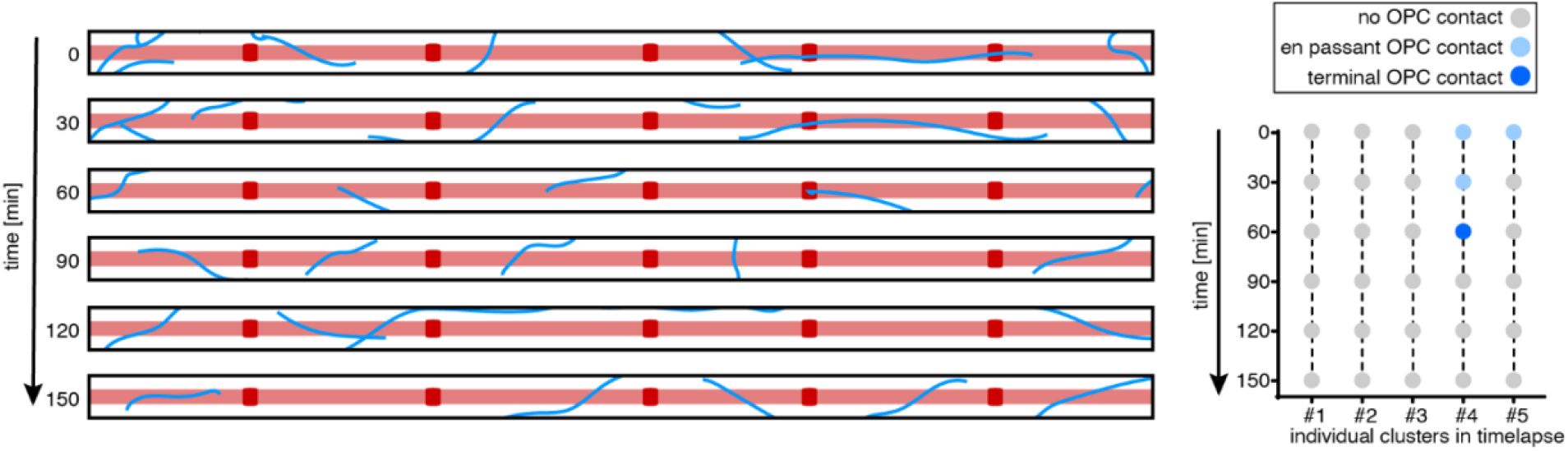
Supplementary to Figure 2. Reconstruction of OPC processes around an unmyelinated axonal stretch containing 5 Nfasca clusters, showing examples of process-cluster interactions and their duration, over time.

**Figure S3:**
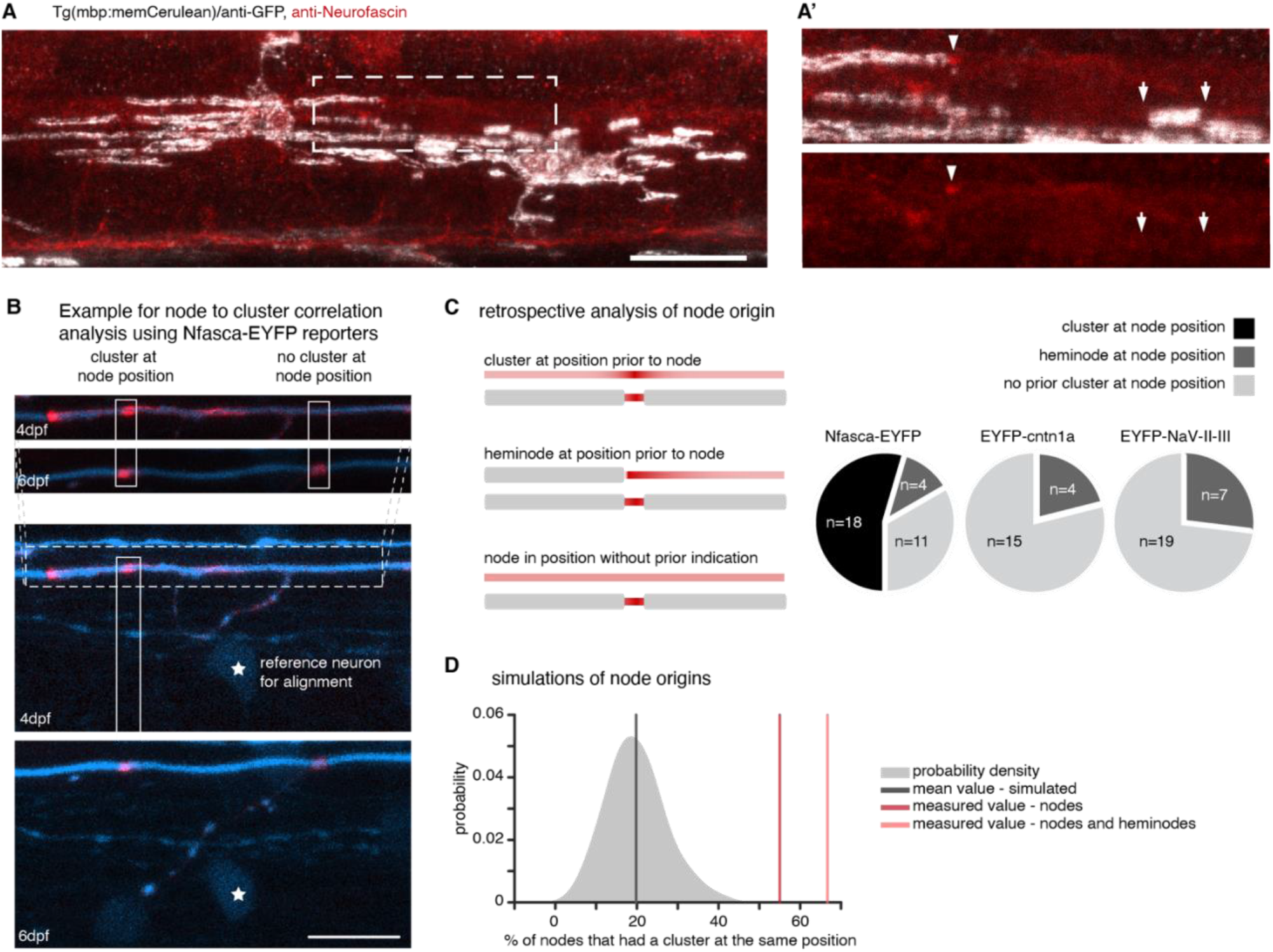
Supplementary to Figure 3. **A)** Confocal images of whole-mount immunohistochemistry against Nfasca (red) in the spinal cord of a full transgenic line with all myelin labelled (+staining against GFP-grey). **A’**) Magnification of the boxed area in panel A. Arrowheads point to Nfasca positive heminodes while arrows point to Nfasca negative heminodes. Scale bar: 20 μm. **B)** Example confocal images of the same axon stretch before and after its myelination. Images from the two timepoints were aligned to a local landmark (asterisk) and the positions of clusters at the earlier timepoint were compared with the position of the future node. Scale bar 10 μm. **C)** Retrospective analysis of node origin. Cartoon showing three possible origins of nodes. (Right) Pie charts show quantification of node origin for Nfasca-EYFP, EYFP-cntn1a and EYFP-NaV-II-III. **D)** Simulation of node origin and their predicted co-localisation with nodes when random positioning is assumed. The shaded curve shows the probability distribution for the simulated values (grey line indicates the mean of simulated data). The light red and dark red lines indicate the quantified cluster fates analysis shown in panel C.

**Figure S4:**
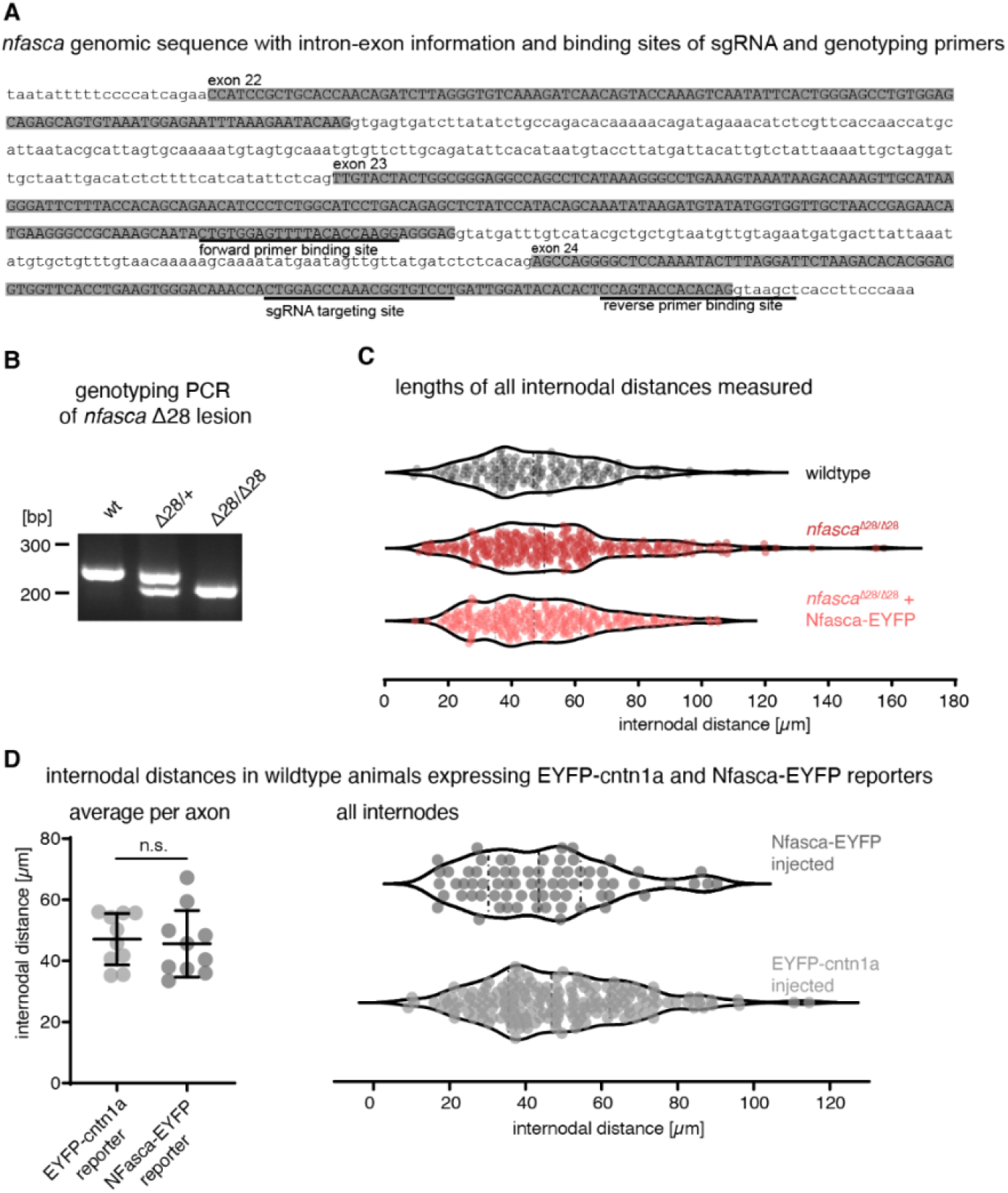
Supplementary to Figure 4. **A)** Part of the *nfasca* genomic sequence around the sgRNA target site. Intron sequences are shown in lowercase, exon sequences are capitalised and highlighted in grey. Genotyping primer sequences and gRNA recognition site are underscored. **B)** Example of genotyping PCR results showing wildtype, heterozygous (Δ28/+) and homozygous (Δ28/Δ28) mutants for the *CRISPR-Nfasca*^Δ28^. **C)** Distribution of all internodal distances quantified in Fig. 3G showing all data points. **D)** Quantification of internodal distances using Nfasca-EYFP and EYFP-cntn1a reporters. (Left) Average internodal distance per axon. Data shown as mean ± SD (unpaired t test). (Right) Distribution of all measured internodal distances.

**Table S1:**
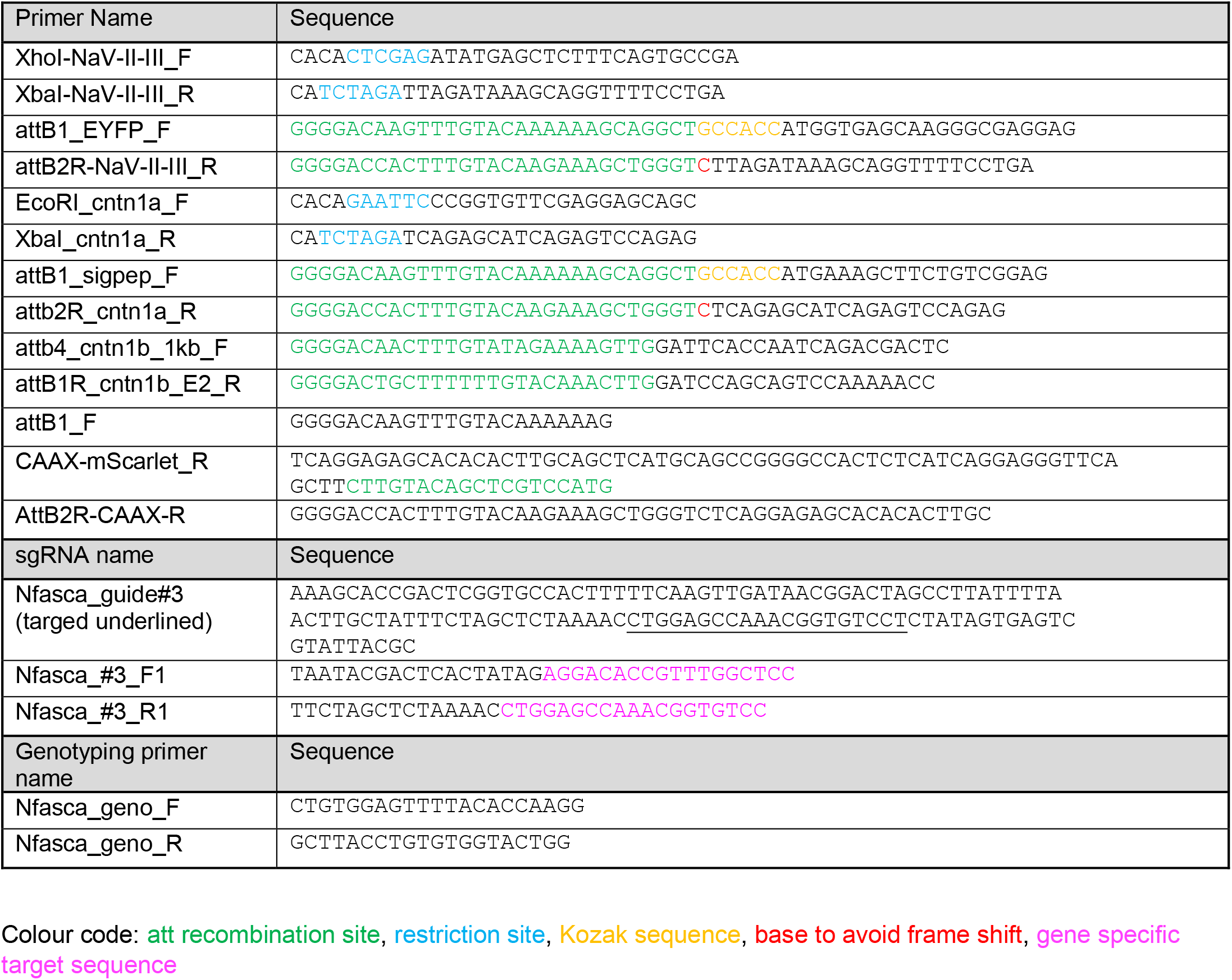
Primers used in this study.

